# Lipid droplets are intracellular mechanical stressors that promote hepatocyte dedifferentiation

**DOI:** 10.1101/2022.08.27.505524

**Authors:** Abigail E. Loneker, Farid Alisafaei, Aayush Kant, Paul A. Janmey, Vivek B. Shenoy, Rebecca G. Wells

## Abstract

Matrix stiffening and external mechanical stress have been linked to disease and cancer development in multiple tissues, including the liver, where cirrhosis (which increases stiffness markedly) is the major risk factor for hepatocellular carcinoma. Patients with non-alcoholic fatty liver disease and lipid-droplet-filled hepatocytes, however, can develop cancer in non-cirrhotic, relatively soft tissue. Here, we show that lipid droplets are intracellular mechanical stressors with similar effects to tissue stiffening, including nuclear deformation, chromatin condensation, and hepatocyte dedifferentiation. Mathematical modelling of lipid droplets as inclusions that have only mechanical interactions with other cellular components generated results consistent with our experiments. These data show that lipid droplets are intracellular sources of mechanical stress and suggest that nuclear membrane tension integrates cell responses to combined internal and external stresses.

**Significance Statement:** Deformation of the nucleus as a result of *extracellular* sources of stress, including increased substrate stiffness, constricted migration, and compression, has been well documented to lead to increased nuclear rupture, changes in gene expression, and accumulation of DNA damage. Lipid droplet accumulation in hepatocytes provides a unique scenario to investigate potential *intracellular* mechanical stresses and sources of nuclear deformation. Our results show that lipid droplets are significant mechanical elements in the cell, deforming the nucleus in a way that promotes hepatocyte dedifferentiation and resisting cytoskeletal contraction and alignment.

## Introduction

Non-alcoholic fatty liver disease (NAFLD), typified by lipid-filled hepatocytes, is the most rapidly increasing liver disease globally (1) and a major precursor of cirrhosis (end-stage liver disease). NAFLD is also an increasing cause of hepatocellular carcinoma (HCC)(2), a primary liver cancer leading to 800,000 deaths worldwide per year (3). Cirrhosis, which is characterized by advanced fibrosis, architectural reorganization of the liver, and, notably, increased liver stiffness (4-7), is the main risk factor for HCC and is associated with over 80% of HCC cases (8). Liver stiffness measurements correlate with occurrence of HCC and patient prognosis, and are used clinically in HCC surveillance (5-7), consistent with a growing body of mechanobiology literature showing that tissue stiffness contributes to malignancy; mechanical stress from various sources, including matrix stiffness, can lead directly to the accumulation of double-stranded DNA breaks, depletion of DNA repair factors, and chromosomal aberrations (9-12). This suggests a potential role for abnormal mechanics in the development of HCC.

A significant percentage (20–63%) of patients with NAFLD-related HCC, however, have non-cirrhotic and relatively soft tissues (13-20). We hypothesized that lipid droplets deform the hepatocyte nucleus and serve as *internal mechanical stressors*, even in a soft environment, mimicking the effects of an externally stiff environment. Normal hepatocytes are stiffness sensitive, spreading and dedifferentiating in response to increasing substrate stiffness (21-23). Recent research also shows that cells in stiff, cirrhotic livers have deformed nuclei (24), suggesting a link between nuclear deformation and hepatocyte dedifferentiation. The nucleus acts as a central integration site for mechanical stress in multiple cell types, sensing deformation through increases in nuclear membrane tension and dynamically adapting to the local environment (25, 26), potentially through persistent changes in gene expression (27, 28). Deformation can alter gene expression epigenetically and through alteration of 3D chromatin structure, changing the accessibility and transcription of relevant genes (29). Furthermore, nuclear deformation contributes to DNA damage accumulation through increased replication stress (12) and nuclear rupture (9-11, 30-32). The cytoskeleton plays an important role in transmitting external stresses to the nucleus, and disruption of nucleo-cytoskeleton links is often sufficient to reverse stiffness-mediated effects (24, 33-35).

We previously showed that lipid droplets disrupt normal mechanosensing in primary human hepatocytes (36). Our goal in this study was to define experimentally and with theory the interactions between lipid droplets, the cytoskeleton, and the nucleus, and to determine their consequences for hepatocyte function. Previous studies of nuclear deformation investigated *extracellular* sources of stress, including increased substrate stiffness, constricted migration, and compression. Lipid droplet accumulation in NAFLD provides a unique scenario to investigate potential *intracellular* mechanical stresses and sources of nuclear deformation.

## Results

For all experiments, primary human hepatocytes (PHH) were seeded sparsely on soft (500 Pa storage modulus) and stiff (10 kPa storage modulus) polyacrylamide gels coated with collagen, which appear functionally equivalent to normal and cirrhotic liver stiffnesses (36-38). Individual cells not in contact with other cells were analyzed in order to assess the impact of stiffness and to isolate cell-substrate interactions. To induce lipid droplet accumulation, media was supplemented with 400 μM oleate, a non-toxic fatty acid that is the most common monounsaturated fatty acid in the human diet and is readily taken up and processed by hepatocytes into lipid droplets. 400 μM is comparable to the serum concentration of lipid in obese patients (39) and is non-lipotoxic (36). Cells treated with oleate accumulate more lipid and form larger lipid droplets than the common fatty acids palmitate or lineolate (36), making oleate the optimal choice for studying the physical effects of large cytoplasmic lipid droplets.

### Lipid-loaded cells spread less and have smaller nuclei

PHH took up a significant amount of oleate when cultured on substrates of all stiffnesses (Fig. S1A, B). Although lipid mean intensity and lipid area fraction were higher in cells on soft compared to stiff substrates (Fig. S1A, B), there were no differences in total lipid volume (Fig. S1c), suggesting that stiffness does not alter total processing of fatty acids. Average lipid droplet size in oleate-treated cells decreased with increasing stiffness (Fig. S1D). Hepatocytes increased their volume and spread more on substrates of increasing stiffness, and this relationship was maintained in oleate-treated cells (Fig. 1A, B). Compared to controls, oleate-treatment reduced cell area on all stiffnesses without altering cell volume, indicating that lipid droplets reduced the ability of cells to spread. The equivalent cell volumes between control and lipid-loaded cells (Fig. 1A) suggest that effective cytoplasmic volume is decreased, potentially impacting molecular crowding, osmotic pressures, and reaction rates. Nuclear cross-sectional area and volume also increased in response to stiffness in all cells; however, lipid-loaded cells had smaller and less spread nuclei than controls on any given stiffness (Fig. 1C, D), which was also reflected in a significant shift downward observed in the relationship between cell and nuclear volume in oleate-treated compared to control cells (Figure 1E). This effect is seen for area as well as volume and is thus not explained by lipid-driven changes in cell spreading (Fig. S1E); however, given the proportional relationship between nuclear and cell volume (which is linearly related to cytoplasmic volume) for a given cell type (40), it is consistent with a decrease in effective cytoplasmic volume. This was confirmed by plotting nuclear volume against estimated cytoplasmic volume (cell volume -nuclear volume - lipid volume), which showed no difference in the observed relationship between control and oleate treated cells (Figure 1F).

**Figure 1.**
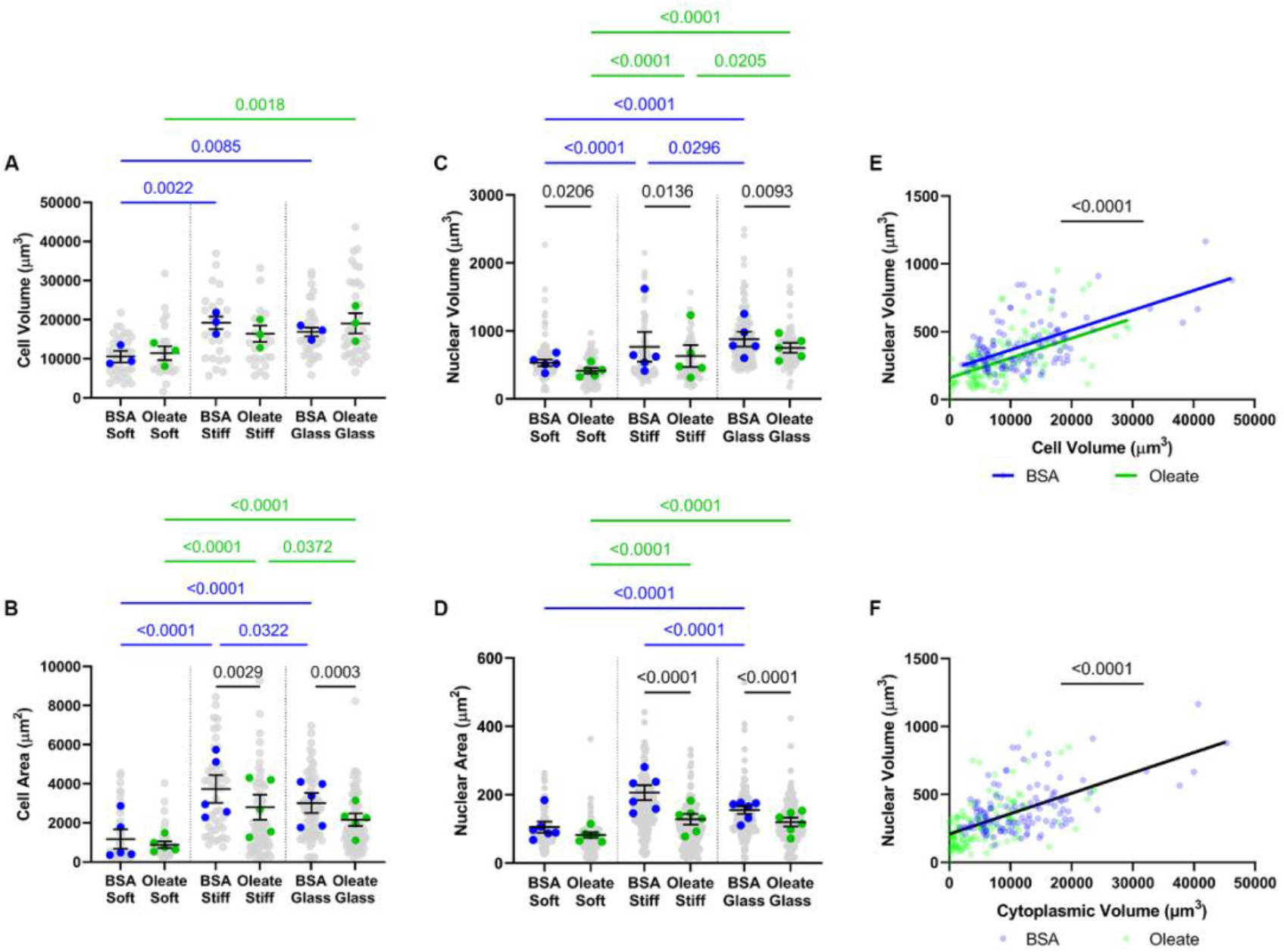
Lipid-droplet accumulation decreases nuclear size and disrupts nuclear/cytoplasmic volume ratios. (**A**) Cell volume (*n = 3*), (**B**) cell area (*n = 5*), (**C**) nuclear volume (*n = 5*), and (**D**) nuclear area (*n = 6*) in control and oleate-treated cells on soft PAA, stiff PAA, and glass. (**E**) Scatter plot comparing cell volume and nuclear volume in individual cells, pooled across stiffnesses, fit with linear regression. (**F**) Scatter plot comparing estimated cytoplasmic volume and nuclear volume in individual cells, pooled across stiffnesses, fit with linear regression. Statistics: (A-D) Data are the mean +-s.e. of *n* independent experiments, p-values calculated using two-way ANOVA. Colored dots are the means per experiment, while the grey dots are the individual cell values. (E, F) Data are the values for individual cells from *n = 3* independent experiments. Blue indicates control cells, while green indicates oleate-treated cells. (E) Slope of both lines is significantly non-zero (p<0.0001 for both control and oleate-treated cells) but not significantly different from one another, so a pooled slope was calculated and used. Y-intercepts are significantly different (p = 0.0017). (F) Linear regression indicated that the slope for both groups was significantly non-zero but not significantly different from one another; neither were the Y-intercepts, so a line was fit to the pooled data.

### Lipid droplets deform hepatocyte nuclei

Lipid droplets deform the nuclear membranes of individual cells on all stiffness substrates (Fig. 2A). While classic shape analysis parameters, including circularity, aspect ratio, roundness, and solidity, showed significant differences between control and oleate-treated cells (Fig. S2), we developed a deformation parameter that was sensitive to and inclusive of both large- and small-scale deformations in order to fully capture nuclear membrane shape changes. We wrote a custom MATLAB program that exploits the circularity of hepatocyte nuclei to quantify areas of local deformation. The program identifies the center point of each nucleus and from this point linearizes the nuclear membrane boundary (measuring from the center point to the nucleus boundary), normalizes to the mean radial distance and plots the radial distance against the angle. From this graph, we calculate the irregularity: the area between the linearized membrane and a line representing the boundary of a perfect circle (Fig. 2B). Control cells on all stiffness substrates had a very low irregularity (Fig. 2C), but irregularity was increased in oleate-treated cells on all stiffnesses, consistent with the nuclear indentation by lipid droplets we observe (Fig. 2A, C). By calculating the local curvature, we also captured the acuteness of indentations in the nuclear membrane (Fig. 2B). Histograms of curvature frequency demonstrated that oleate-treated cells on all stiffnesses have a higher frequency of sharp indentations (of both positive and negative curvature) with sub-micron radii (Fig. 2E). The distribution of indentations for control cells is symmetric across the y-axis, consistent with random membrane fluctuations, while the distribution for oleate-treated cells is not, further confirming that these sharp indentations are imposed locally on the nuclear membrane (Fig. S3). Interestingly, there seems to be a higher frequency of high *positive* curvature membrane regions than negative curvature regions in oleate-treated cells, suggesting that high curvature is more likely to occur indirectly than as a result of direct indentation by high curvature lipid droplets (Fig. S3). In contrast to the changes in nuclear irregularity, aspect ratio metrics in the YZ plane showed that the nuclei of oleate-treated cells remained similarly rounded on all stiffness substrates and did not exhibit the stiffness-dependent flattening (increased aspect ratio) seen in controls (Fig. 2D). Thus, as the oleate-treated cells became more spread, lipid droplets in the cytoplasm resisted nuclear deformation by cortical actin and stress fibers. This suggests that lipid droplets have dual mechanical roles in the cell: while indenting the nuclei radially, they prevent deformation in the z-direction caused by cell spreading.

**Figure 2.**
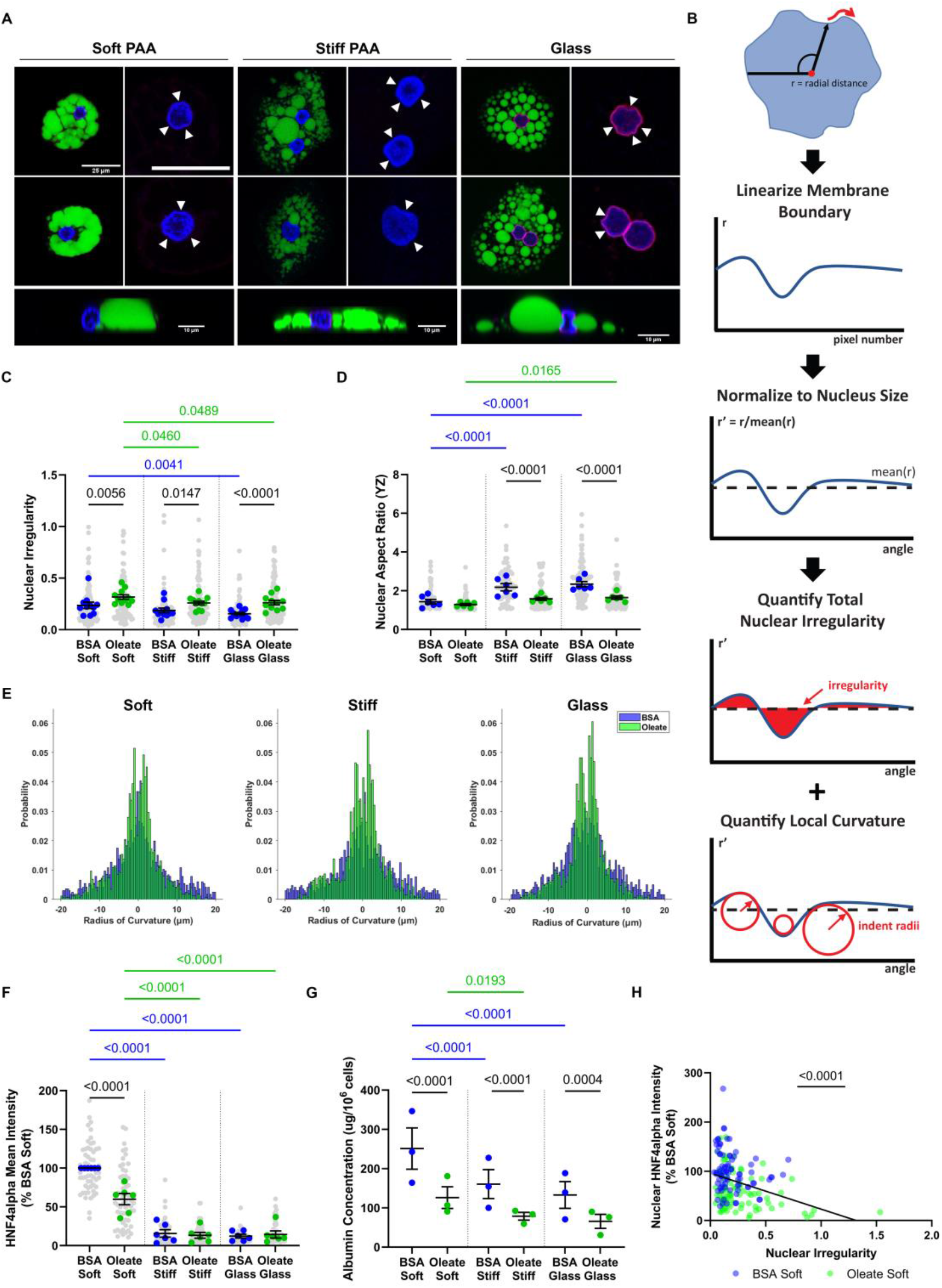
Lipid droplets indent hepatocyte nuclei, leading to dedifferentiation. (**A**) Nuclei of oleate-treated cells are indented by lipid droplets. Under each heading for stiffness, left column images show representative cells with lipid droplets and the right column images are zoomed-in views of the same cell nucleus with white arrowheads used to emphasize areas of indentation. Bottom row shows examples of deformation in the YZ plane. Scale is the same for first two rows of images, with the bar = 25 μm. YZ-cross sections have individual scale bars representing 10 μm. DAPI (blue), BODIPY (green), lamin A/C (red). (**B**) Schematic representation of nuclear irregularity, the parameter used to quantify nuclear deformation in the XY plane. (**C**) Nuclear irregularity (*n = 5*) and (**D**) YZ aspect ratio (*n = 3*) in control and oleate-treated cells. (**E**) Histograms of the radii of curvature of nuclear membrane indentations. (**F**) HNF4α mean intensity (*n = 6*, normalized to control on soft) in the nuclei of control and oleate-treated cells. (**G**) Albumin concentration (*n = 3*) in the media of control and oleate-treated cells. (**H**) Scatter plot with linear regression of nuclear irregularity vs. nuclear HNF4α intensity in individual cells on soft substrates. Statistics: (C, D, F, G) Data are the mean +-s.e. of *n* independent experiments. P-values were calculated using two-way ANOVA with multiple comparisons. (E) Data are the individual dent radii pooled across cells from n = 3 independent experiments. On each stiffness, control and oleate-treated distributions compared with K-S test. (H) Data are the values for individual cells on soft gels from n = 3 independent experiments. P-value calculated with an F-test. In all cases blue indicates control cells, while green indicates oleate-treated cells.

### Lipid droplets promote hepatocyte dedifferentiation

Hepatocyte differentiation is stiffness sensitive, with optimal hepatocyte function occurring when cells are plated on soft substrates (23). Given that stiffness deforms the nuclei of liver cells and that this deformation mediates stiffness-driven phenotypic effects (24), we studied whether the deformation of nuclei by lipid-droplets similarly induces hepatocyte dedifferentiation. In control and in oleate-treated cells, albumin production and nuclear HNF4α intensity decreased significantly with increasing substrate stiffness (Fig. 2F, G), as predicted from the literature, but nuclear HNF4α was also significantly decreased in oleate-treated cells on soft substrates (Fig. 2F). Expression of HNF4α was negatively correlated with nuclear irregularity on soft substrates, and the relationship was similar for control and lipid-loaded cells (Fig. 2H). Lipid-loaded cells also secreted less albumin than controls (Fig. 2G). This suggests that on soft substrates, radial deformation of the nucleus by lipid droplets promotes hepatocyte dedifferentiation and shows that the presence of lipid droplets has similar effects as stiff substrates.

HNF4α is a key transcriptional regulator in hepatocytes that preferentially associates with high-mobility, decondensed chromatin (41), such that increases in chromatin condensation could decrease expression levels. Therefore, we used image analysis of DAPI-stained nuclei(42) to determine the chromatin condensation state of oleate-treated cells, observing that chromatin condensation increases with both substrate stiffness and lipid-loading (Fig. 3A). Chromatin distribution between expressive euchromatin and compacted heterochromatin phases has been proposed to be a result of liquid-liquid phase separation of chromatin(43). However, the role of mechano-osmotic forces in regulating the chromatin reorganizational response of the nucleus is not yet fully understood. To this end, we developed a thermodynamically consistent phase-field model (see Methods) to predict the spatial distribution of chromatin in the nucleus due to either extra-nuclear mechano-osmotic loads, or mechano-chemical cues in the form of substrate (or extracellular matrix) stiffness (29).

**Figure 3.**
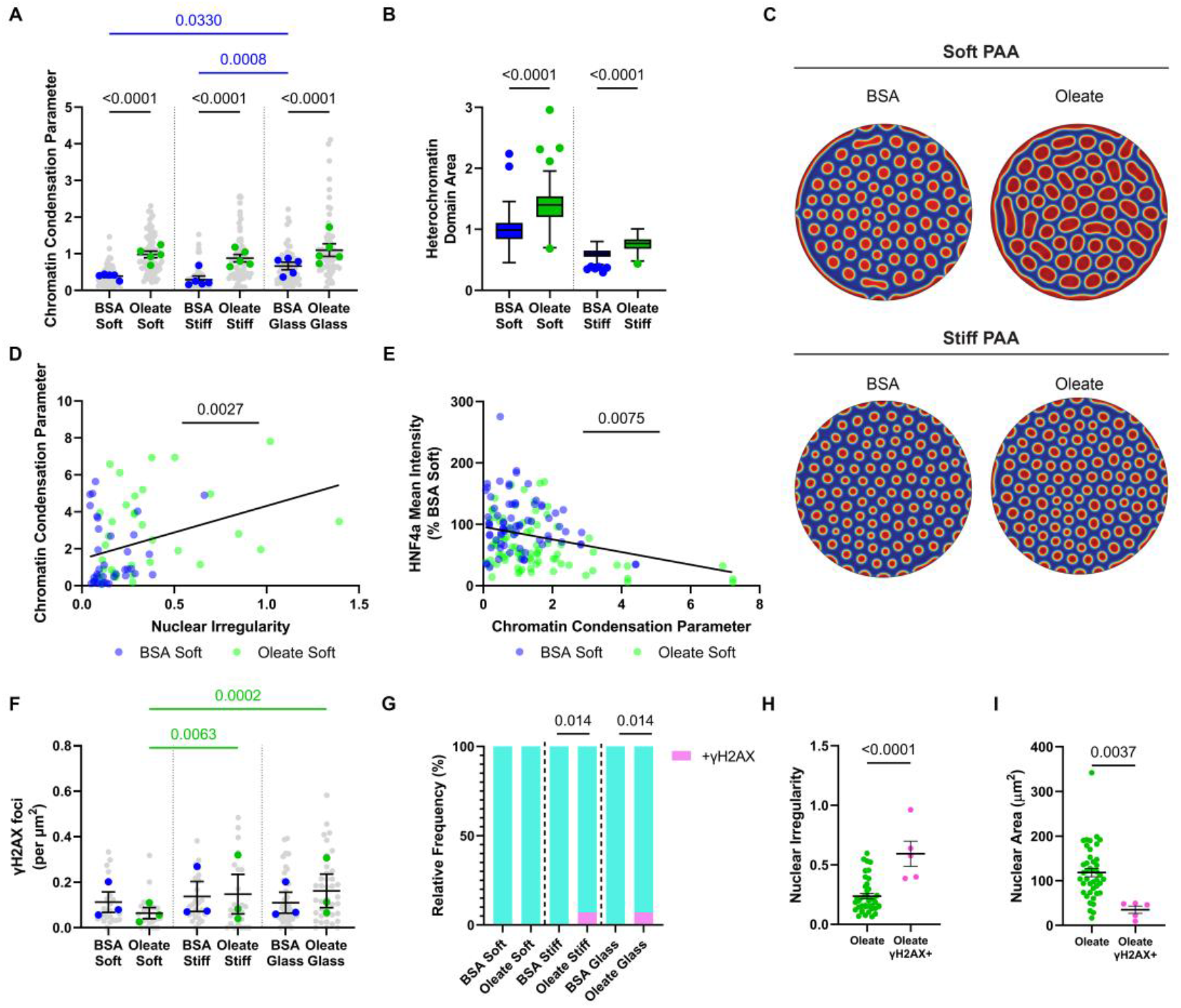
Lipid loading condenses chromatin and can promote DNA damage. (**A**) Chromatin condensation state (*n = 5*) of control and oleate-treated cells. (**B**) Quantification and (**C**) visualization of modelled chromatin phase separation in response to mechano-osmotic forces. Circular images represent heterochromatin domain size and organization within simulated cell nuclei for control and oleate-treated cells (red representing compact heterochromatin, blue uncompact euchromatin). The average size of heterochromatin domains over repeated simulations is quantified in B. (**D**) Scatter plot with linear regression of chromatin condensation parameter vs. nuclear HNF4α intensity in individual cells on soft substrates. (**E**) Scatter plot with linear regression of nuclear irregularity vs. chromatin condensation parameter in individual cells on soft substrates. (F) γH2AX foci *(n=3*) normalized to nuclear cross-sectional area in control and oleate-treated cells. (**G**) Percentage of cells staining fully positive for γH2AX. (**H**) Nuclear irregularity and (**I**) cross-sectional area of oleate-treated cells depending on γH2AX staining. Statistics: (A, F) Data are the mean +-s.e. of *n* independent experiments. P-values were calculated using two-way ANOVA with multiple comparisons. (D, E) Data are the values for individual cells from *n = 3* independent experiments. P-value calculated with an F-test. (G) Data are the percentage of total cells from *n = 3* independent experiments. P-values calculated with Chi-squared test comparing control to oleate-treated distribution on each stiffness. (H, I) Data points represent individual cells from the two experiments where fully positive γH2AX nuclei were observed. This subset of cells was compared to the others with an un-paired T-test. In all cases blue indicates control cells, while green indicates oleate-treated cells.

The model incorporates the following key features – (i) energetics of chromatin-chromatin interactions enabling the separation of dissimilar phases, (ii) chromatin-lamina interactions enabling formation of lamina-associated domains, (iii) diffusion kinetics of nucleoplasm and chromatin enabling a quantity-conserving flow of chromatin, and (iv) non-conservative reaction kinetics capturing the methylation or acetylation of chromatin via epigenetic regulation pathways, allowing an interconversion of heterochromatin and euchromatin phases (44). Consistent with our experimental results showing decreased nuclear volume (Fig. 1C), we can simulate the presence of compressive forces on the nuclei due to lipid droplets via the mechano-osmotic compression of the nuclei, driving an outflux of water from the nucleoplasm into the cytoplasm. By allowing a non-zero flux along the periphery of the simulated nucleus, we allow the diffusive exchange of water between the nucleus and the cytoplasm driven by changes in its chemical potential (Eq. 2, Methods). The model predicts that oleate-treated cells exhibit increased chromatin compaction on both soft and stiff substrates, increasing the area of compacted heterochromatin domains (Fig. 3B-C), consistent with experimental results (Fig. 3A).

Chromatin condensation and nuclear irregularity were negatively correlated with HNF4α intensity in oleate-treated cells on soft substrates, and positively correlated with each other (Fig. 3D, E). This combined with the modeling supports a potential mechanism by which lipid accumulation induces nuclear deformation and decreases nuclear volume, leading to condensed chromatin and reduced expression of HNF4α and downstream hepatocyte-specific genes.

### A small subset of lipid-loaded cells have increased DNA damage

DNA damage can be induced by nuclear deformation in migrating cancer cells (12). We stained with γH2AX to determine the frequency of double-stranded DNA breaks in lipid-loaded cells with nuclear deformation. There was no overall difference in the frequency of double stranded breaks in control and oleate-treated cells (Fig. 3F). However, a subset of oleate-treated cells on stiff gels and glass had a large number of γH2AX-positive foci (Fig. 3G). Interestingly, these nuclei were more irregular and had smaller cross-sectional area than other oleate-treated cells (Fig. 3H, I), suggesting that compression and indentation by droplets increases the likelihood of DNA damage, but only in extreme cases.

### Lipid droplets disrupt cytoskeletal networks

A previously published chemo-mechanical model of cytoskeletal-nuclear mechanosensing proposes that actin and microtubules act in opposition to determine the flattening and deformation of the nucleus and to generate changes in mechanosensitive signaling (44). We therefore evaluated actin and microtubule networks in lipid droplet-laden hepatocytes.

Control cells on soft substrates exhibited high intensity phalloidin staining, showing actin that was disorganized and diffuse and generally localized to the basal cell membrane (Fig. 4A, B, Fig. S4A). With increasing stiffness, phalloidin intensity (indicative of concentration) and integrated density (indicative of total content) decreased in control hepatocytes and actin fibers became distinct, moving to the apical cell membrane (Fig. 4A, B, Fig. S4A, B). In oleate-treated cells, actin intensity and integrated density were decreased on soft substrates (Fig. 4B, Fig. S4B), suggesting that stiffness and lipid loading have similar effects on actin in hepatocytes. Actin fibers were displaced by lipid droplets, particularly in areas with large droplets (Fig. 4A). Organization was quantified by assessing fiber length and junction density. Control cells demonstrated a stiffness-dependent increase in fiber length and decrease in junction density, consistent with the appearance of stress fibers (Fig. 4C, D). Oleate-treated cells had consistent fiber length and junction density without stiffness-related changes, but were significantly different than control cells: they had longer fibers and lower junction density than control cells on soft substrates and shorter actin fibers and higher junction density on stiff substrates (Fig. 4C, D). We used vector analysis to determine the orientation distribution of actin fibers and to classify cells based on their level of actin fiber alignment. For all stiffnesses, oleate-treated cells had lower levels of alignment than controls, with more cells having multiple fiber directions (Fig. 4E, note increase in unaligned (teal) and decrease in single alignment direction (pink)). Thus, lipid droplets impose a disorganized and branched actin network structure on soft substrates and prevent stiffness-driven fiber alignment.

**Figure 4.**
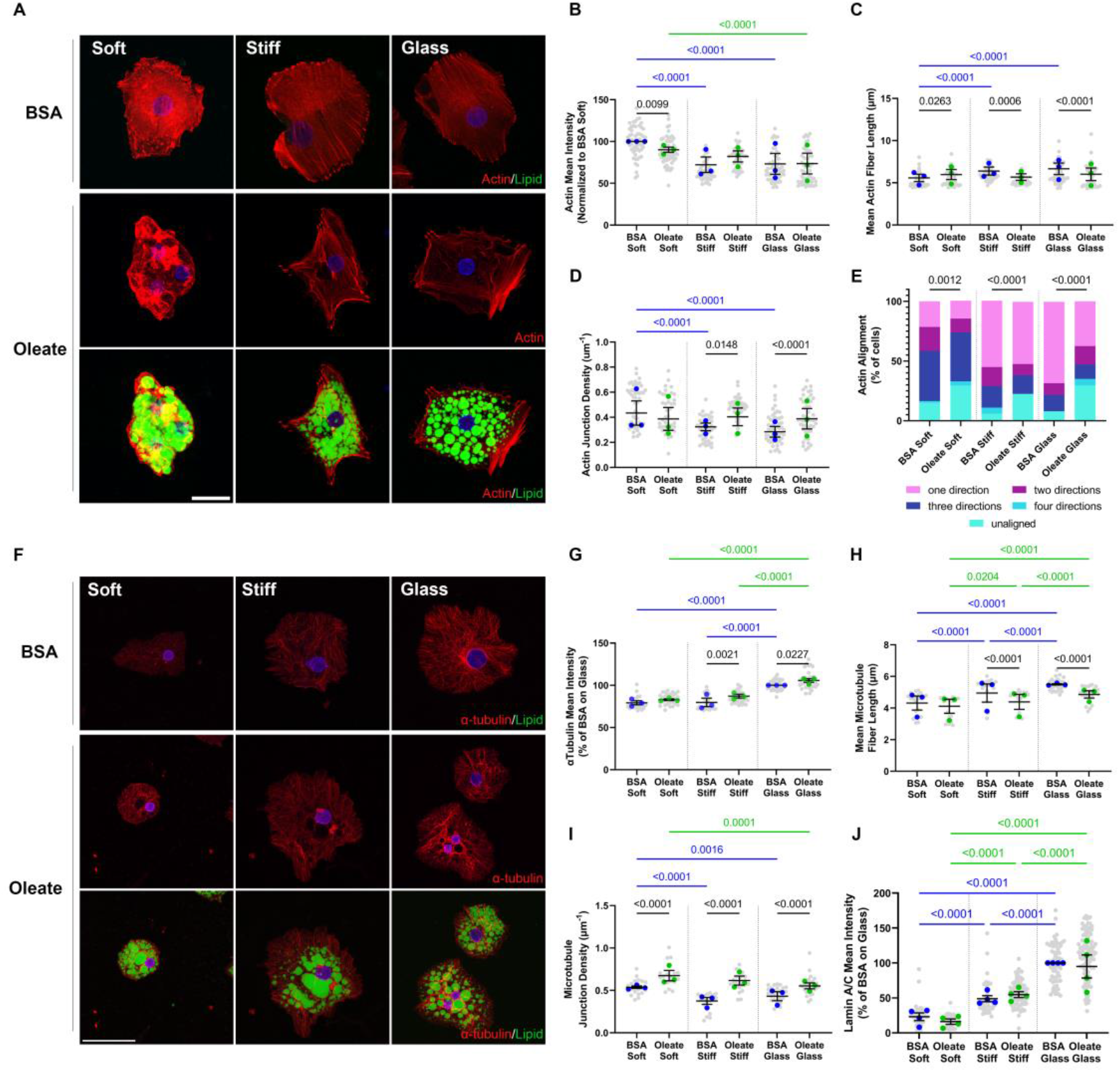
Lipid droplets disrupt cytoskeletal fibers. (**A**) Representative images of actin organization (phalloidin) in control and oleate-treated cells on soft PAA, stiff PAA, and glass. Oleate-treated cells are shown with (bottom row) and without (middle row) lipid droplets. Scale bar is 25 μm and is the same for all images. DAPI (blue), BODIPY (green), phalloidin (red). (**B**) Mean phalloidin intensity (normalized to mean of control on soft) for control and oleate-treated cells. (**C)** Mean fiber length and (**D**) junction density of actin fibers detected by image analysis of maximum z-projections. (**E**) Frequency of aligned actin fiber directions in control and oleate-treated cells. Cells with more than one alignment direction indicate groups of fibers aligned in different directions within a single cell. (**F**) Representative images of microtubule organization (α-tubulin) in control and oleate-treated cells. Oleate-treated cells are shown with (bottom row) and without (middle row) lipid droplets. Images are maximum Z-projections. Scale bar is 50 μm and applies to all images. DAPI (blue), BODIPY (green), and α-tubulin (red). (**G**) Mean α-tubulin intensity (normalized to mean of control on glass) for control and oleate-treated cells. (**H**) Mean fiber length and (**I**) junction density of microtubules detected by image analysis of maximum Z-projections. (**J**) Mean lamin A/C staining (normalized to mean of control on glass) for control and oleate-treated cells. Statistics: (B-D, G-I). Data are the mean +-s.e. of n = 3 independent experiments. (E) Data are the percentage of total cells from n = 3 independent experiments. P-values were calculated with Chi-squared test comparing control to oleate-treated distribution on each stiffness. (J) Data are the mean +-s.e. of *n = 4* independent experiments. P-values were calculated using two-way ANOVA with multiple comparisons. In all cases, blue indicates control cells, green indicates oleate-treated cells.

Lipid droplets also disrupted microtubules. In both control and oleate-treated cells, α-tubulin intensity and integrated density increased with increasing substrate stiffness (Fig. 4F, G, Fig. S4C). On all stiffnesses and with oleate treatment, microtubules were distributed throughout the cytoplasm (Fig. S4D). However, oleate-treated cells had a higher mean α-tubulin intensity than controls, indicating a higher microtubule density (Fig. 4G). 3D image reconstruction shows microtubules appearing to form cages around individual lipid droplets. This is supported by fiber network analysis, which shows shorter microtubule length and increased branching frequency in lipid-loaded compared to control cells (Fig. 4H, I).

Connections between the cytoskeleton and nucleoskeleton are vital to transmitting external forces to the nucleus. Both control and oleate-treated cells exhibited increasing lamin A/C intensity in response to increased substrate stiffness, as expected with increasing nuclear stress. Surprisingly, despite significant cytoskeletal disorganization and disruption, lamin A/C intensity and localization to the nuclear membrane were not affected by lipid-loading (Fig. 4J). This suggests that the nuclei of oleate-treated cells remain under increased mechanical stress, even as cell-spreading and nuclear compression by stress fibers are reduced.

### Lipid droplets reduce traction forces

To further investigate the role lipid droplets play in resisting stiffness-driven changes to the cytoskeleton, we measured hepatocyte-generated traction forces and found that lipid-loading significantly decreased the mean and maximum traction force (Fig. 5A, B). While both control and oleate-treated cells showed a positive correlation between cell area and force generation, the relationship was significantly different for oleate-treated cells (Fig. 5C, Fig. S5A) – indicating that reductions in traction force are not simply due to a lipid-driven decrease in cell spread area. Furthermore, the force generated in oleate-loaded cells is negatively correlated to lipid density (Fig. 5D, Fig. S5B), suggesting that lipid-droplets resist cell-generated contractility. These results indicate that lipid accumulation disrupts hepatocyte cytoskeletal networks, rendering them both less able to generate traction forces and less stiffness sensitive than controls.

**Figure 5.**
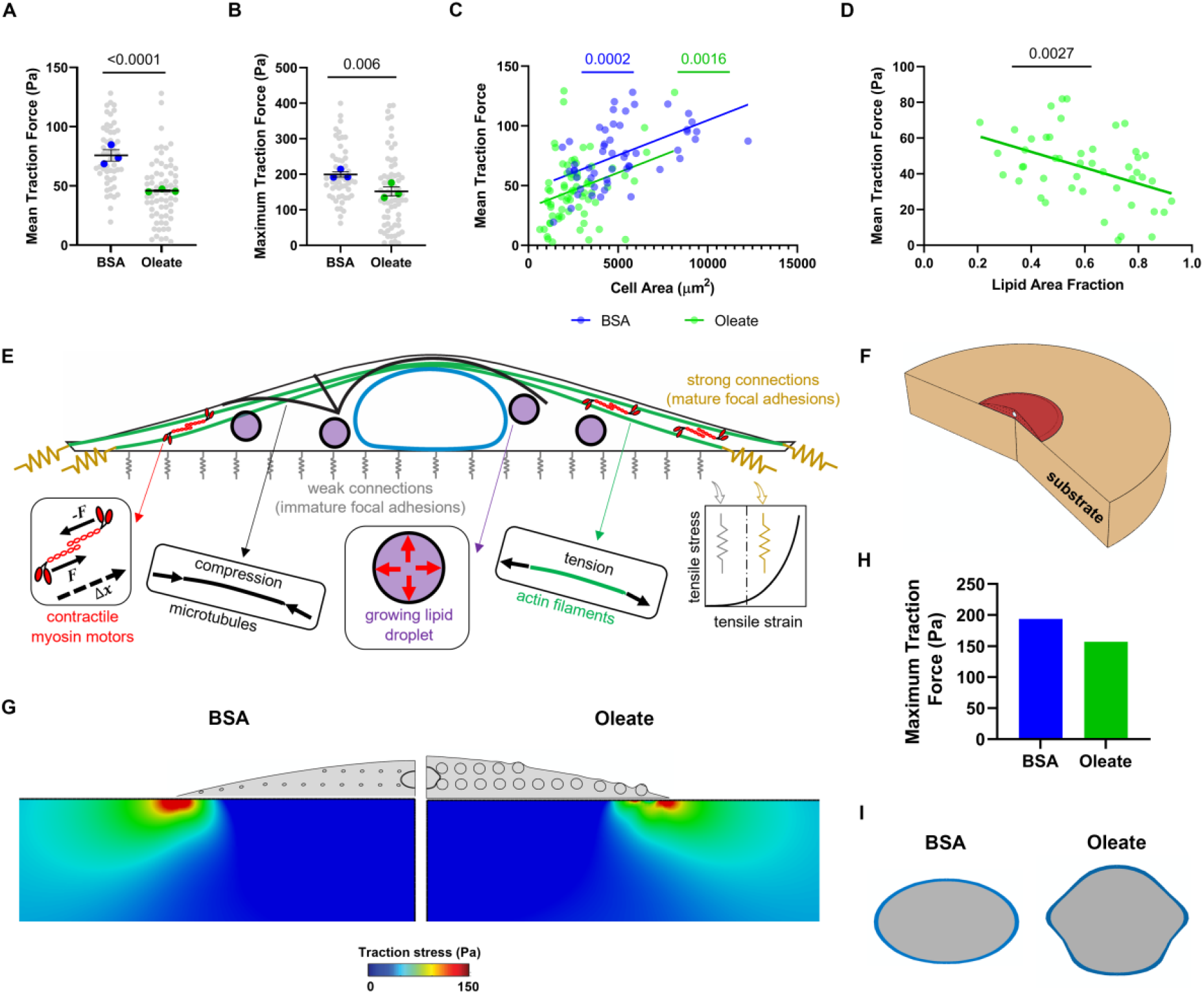
Experimental and modeling approaches show that lipid droplets reduce cell force generation. (**A**) Mean and (**B**) maximum traction forces for control and oleate-treated cells on stiff PAA gels. (**C**) Scatter plot with linear regression of cell area versus mean traction force in individual cells for control and oleate-treated cells. (**D**) Scatter plot and linear regression of lipid droplet density and mean traction force. (**E**) Schematic of chemo-mechanical model of lipid-droplet cytoskeletal interactions in the generation of traction forces. Adapts previously published model(48) and adds lipid droplet growth in the cytoplasm. (**F**) Geometry of the cell and substrate over which cell traction forces were modelled, red indicating the cell and tan indicating the substrate. (**G**) Visualization and (**H**) quantification of reduced traction forces in lipid-loaded cells as predicted by our mathematical model. (**I**) Visualization of lipid-droplet associated nuclear deformation as predicted by our mathematical model. Statistics: (A, B) Data are the mean +-s.e. of *n = 3* independent experiments. P-values were calculated using two-way ANOVA with multiple comparisons. (C, D) Data are individual cells from *n = 3* independent experiments. P-values calculated with F-test. In all cases, blue indicates control cells, while green indicates oleate-treated cells.

### Mathematical modelling suggests mechanical interaction between droplets and other cellular components is sufficient for nuclear indentation and reduction in traction forces

The previously mentioned chemo-mechanical model of the cytoskeleton (44) was modified to add lipid droplets, to determine whether decreases in traction force could be explained by their physical presence (Fig. 5E). The three-dimensional cell model includes the following components: the cytoskeleton, the focal adhesions, and the nucleus (see reference 44 for details). Lipid droplets were modelled as spherical inclusions that include a thin membrane with internal pressure representing the enclosed fluid. The internal pressure was simulated by applying uniform and outward force spatially perpendicular to the internal surface of the membrane which in turn generates tangential tensile stresses in the membrane representing the surface tension in lipid droplets. The inclusions were arrayed within the cell cytoplasm of circular, well-spread hepatocytes (Fig. 5E, F). Note that the droplets can only have mechanical interactions with other cellular components as described in Materials and Methods. For cells with the same area, the addition of lipid droplets reduced the magnitude of traction force that the cells were able to generate, in strong agreement with the experimental results (Fig. 5G, H). Furthermore, the presence of these growing droplets was sufficient to indent the cell nuclei, further supporting the idea that lipid droplets are a mechanical stress on the nucleus (Fig. 5I).

### Altering contractility modulates lipid-associated hepatocyte dedifferentiation

We treated lipid-loaded cells with cytoskeletal inhibitors to test whether the deleterious effects of lipid droplets result from active cytoskeletal forces pushing them into the nucleus. Treatment with blebbistatin, which disrupts myosin, and latrunculin A, which disrupts actin, led to decreases in nuclear area and nuclear aspect ratio in control cells on stiff substrates, as expected given that actin-mediated spreading is inhibited (Fig. 6A, B, Fig. S6A, B). Nocodazole, which depolymerizes microtubules, led to increases in nuclear area and aspect ratio: as expected given that microtubule depolymerization has been shown to increase contractility(45) (Fig. 6A, B, Fig. S6A, B). However, the impact of cytoskeletal disruption was blunted in control cells on soft and oleate-treated cells on all stiffnesses, such that no drug significantly altered nuclear area or aspect ratio on the PAA gels (Fig. 6A, B, Fig. S6A, B). Even on glass, where blebbistatin decreased area and nocodazole increased aspect ratio in oleate-treated cells, those changes were significantly less than drug-treated control cells (Fig. S6A, B). This is consistent with reduced actomyosin contraction in oleate-treated cells – actin is already disrupted, and traction forces already reduced by droplets and the cytoskeleton is thus less sensitive to further pharmacological alteration. In the case of nocodazole, lipid droplets also resist increases in contractility. Surprisingly, nuclear irregularity was not reduced by inhibiting contraction on stiff substrates (Fig. 6C, Fig. S6C), suggesting that contraction is not required for nuclear indentation by lipid droplets. It is also possible that nuclear membrane tension is reduced in drug-treated cells, making it more prone to fluctuation and deformation.

**Figure 6.**
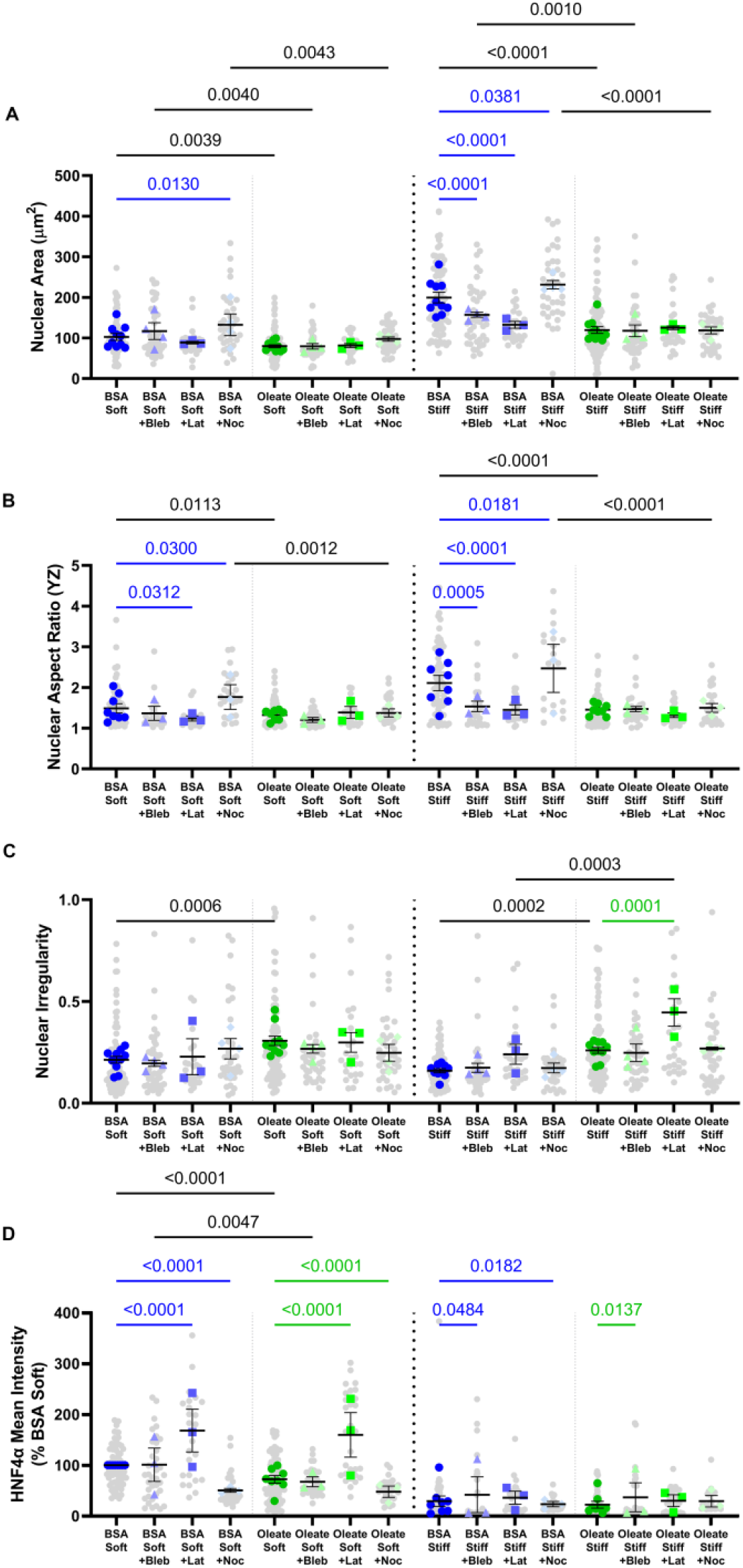
Altering contractility modulates HNF4α expression. (**A**) Nuclear area, (**B**) cross-sectional nuclear aspect ratio, (**C**) nuclear irregularity, and (**D**) mean HNF4α intensity (normalized to control on soft) in control and oleate-treated cells with or without the addition of cytoskeletal drugs on soft and stiff PAA. Blebbistatin (Bleb), Latrunculin A (Lat), Nocodazole (Noc). Data are the mean +-s.e. of *n = 3* for each drug treatment and *n = 8* for the non-treated. P -values were calculated using two-way ANOVA that applied a main effects model with multiple comparisons. Colored significance values indicate changes due to cytoskeletal drugs, while black significance values are due to lipid-loading within a drug-treatment group.

Despite not significantly altering nuclear irregularity, latrunculin-treatment restored HNF4α expression in both control and oleate-treated cells on soft gels, and blebbistatin-treatment partially rescued HNF4α expression in both control and oleate-treated cells on stiff gels (Fig. 6D, Fig. S6D). Nocodazole treatment had the opposite effect, leading to decreased HNF4α expression in both control and oleate-treated cells on soft gels and in control cells on stiff. This shows that interactions between lipid droplets and the nuclear envelope are more deleterious in contractile cells.

## Discussion

We show here that lipid droplets directly indent the hepatocyte nucleus and resist cytoskeletal contraction and that this results in increased chromatin condensation, decreased HNF4α expression, and decreased albumin secretion, consistent with hepatocyte dedifferentiation. Our experimental results were also closely matched by two mathematical models that focused solely on the presence of lipid droplets as incompressible mechanical elements. This shows that lipid droplets are important mechanical elements in the cell, acting as intracellular stressors with effects akin to those of extracellular mechanical stressors like matrix stiffness.

The ability of lipid droplets to deform the nucleus is counterintuitive, given that they are most simply described as an oil-in-water (lipid-in-cytosol) emulsion stabilized by a surfactant (a phospholipid monolayer) (37). The phospholipids, however, create an energy barrier to local surface deformations by increasing the bending energy (37). Additionally, lipid droplet stiffness is a product of the surface tension, which cannot relax without volume changes, while the nucleus is viscoelastic on long time scales. Thus, even though the nucleus is the stiffest organelle in the cell(38), the energy cost of nuclear deformation is less than that of lipid droplet deformation. The magnitude of this barrier can be altered by the phospholipid composition: longer acyl chains and higher membrane concentrations result in a higher energy barrier to deformation. Additionally, different membrane phospholipids have an inherent curvature, and enrichment with negatively curved lipids, such as cholesterol, can promote the fusion of small droplets into larger droplets, which we have seen to be more deleterious (36). This is a potential mechanism for interaction between the biochemical and physical effects of lipid accumulation and suggests that lipid droplet composition may modulate the degree of nuclear disruption. Modelling lipid droplets as growing incompressible spheres recapitulated the nuclear deformation we see in vitro, indicating that they can exert direct force on the nucleus.

The nucleus is an important mechanosensor, both in mediating short term responses to mechanical stimuli and in encoding a long-term “mechanical memory”. Almost all research on the impact of mechanics on the nucleus has used extracellular sources of mechanical stress. Important findings have included the demonstrations that tension exerted on the nuclear membrane can dilate nuclear pores, leading to the translocation of mechanosensitive transcription factors and coactivators, including YAP (33); that cells sense deformation through increases in nuclear membrane tension and dynamically adapt in constricted environments and that there is a “tolerable” level of nuclear deformation, set by the mechanical properties of the cell nucleus(25, 26); and that extreme or persistent levels of nuclear deformation contribute to disease, increasing instances of DNA damage accumulation (9-12, 31). Extracellular sources of mechanical stress used in these studies have included increased tissue stiffness, migration through dense matrices, and the direct application of compressive force. Here we have shown that cells respond to internal sources of mechanical stress in a similar manner, and we have identified nuclear deformation as a potential integration point for the response to combined internal and external forces. This is especially interesting given the disruption and reorganization of the cytoskeletal network caused by lipid loading. Despite reducing cytoskeletal tension, lipid droplets indent nuclei directly, potentially maintaining high nuclear membrane tension even as force transmission through the cytoskeleton is disrupted. This is consistent with our previous work showing that large lipid droplets, which induced larger amounts of nuclear irregularity, increase nuclear translocation of YAP both in vitro and in human samples (36).

Our work has important implications for understanding hepatocyte function in NAFLD, particularly the demonstration that nuclear indentation by lipid droplets on soft substrates reduces expression of HNF4α in a manner proportional to the magnitude of deformation. Lipid-laden hepatocytes are partially resistant to stiffness-driven morphological changes (cell spreading and nuclear flattening) and their nuclei are more resistant to cytoskeletal disruption than control cells. This is because lipid droplets resist cell traction forces, disrupting stiffness-driven alignment of actin fibers and effectively shielding the nucleus from contractile forces. HNF4α expression can be further modulated by altering the contractility of cells with cytoskeletal drugs. On soft substrates, decreasing contractility with latrunculin increased HNF4α in control and oleate-treated cells, fully reversing any reductions due to lipid loading, while increasing contractility with nocodazole did the opposite. Treatment with cytoskeletal drugs also eliminated any correlation between nuclear irregularity and HNF4α expression, suggesting that nuclear deformation is only deleterious when it increases nuclear membrane tension.

Importantly, we saw an increase in double stranded DNA breaks (as indicated by γH2AX) in only a few lipid-loaded cells, suggesting that short-term deformation by lipid droplets is a less severe mechanical stress on the nucleus than external confinement. This may be related to the higher strain rate of migration-associated nuclear deformation(39); while lipid droplets take days to accumulate in culture, a cell can migrate through a constriction in minutes. Lipid accumulation alone may not be sufficient to increase DNA damage but may sensitize nuclei to damage from external mechanical stresses such as increased tissue stiffness. This is consistent with the clinical progression of NAFLD: while NAFLD patients are at a significantly increased *relative* risk of HCC, their absolute risk remains low, lower than would be expected if the cells were regularly incurring significant DNA damage. In our experiments, hepatocytes were only loaded with lipid for 48 hours, while an intact fatty liver would see lipid accumulation and persistence over many years. Lipid accumulation is also dynamic, so human hepatocytes may be subjected to repeated stresses as lipid levels wax and wane, such that even a small increased risk of damage could lead to cancer over a long enough time.

More broadly, this research suggests that stiff intracellular inclusions could be sources of mechanical stress in disease. Glycogen and lysosomal storage diseases affect multiple organ systems including liver, skeletal muscle, heart and brain, and lead to the accumulation of large cytoplasmic inclusions. It is therefore possible that nuclear deformation from intracellular stresses may contribute to disease progression in these cases as well. Overall, our results and others stress the importance of looking at changing mechanical stresses in disease, at both the cell and tissue level, as they are often characterized by multiple and competing disruptions. How cells respond to combination forces, both internal and external, is currently not well understood and is an area ripe for additional research.

## Materials and Methods

### Polyacrylamide Gel Preparation

Polyacrylamide (PAA) gels with a storage modulus of 500 Pa or 10 kPa were made as previously described(40). Cells were also cultured on glass (∼GPa) as a non-physiologically stiff control. Glass coverslips and PAA gels activated with sulfo-SANPAH (Thermo Fisher Scientific, Waltham, MA) were incubated with 0.1 mg/mL rat tail collagen type 1 (Corning, Edison, NJ) for 2 h at room temperature to coat the surface with extracellular matrix ligands and enable cell adhesion. Gels were sterilized by UV exposure for 30 min before culture.

### Cell Culture

Cryo-preserved primary human hepatocytes (PHH) from single donors (BD Gentest, Tewksbury, MA; Lonza, Walkersville, MD; and Thermo Fisher Scientific) were used for all in vitro experiments. PHH were thawed according to the supplier’s instructions using Cryopreserved Hepatocyte Recovery Medium (CM7000; Thermo Fisher Scientific), resuspended in Williams Medium E (MilliporeSigma, Burlington, MA) with plating supplements (CM3000; Thermo Fisher Scientific) and plated at a density of 50,000 cells/cm^2^.

One day after seeding, PHH were serum starved overnight in serum-free Williams Medium E with maintenance supplements (CM4000; Thermo Fischer Scientific). Cells were then incubated for 48 h in DMEM supplemented with 0.5% bovine serum albumin (BSA, free fatty acid (FFA) free; MilliporeSigma) and 1% penicillin-streptomycin with or without the addition of 400 μM sodium oleate. To solubilize the fatty acids and facilitate uptake by PHHs, sodium oleate was preconjugated to BSA. 20 mM oleic acid solution was prepared in 0.01 M NaOH and incubated for 30 min at 70°C. Next, the solution was diluted to 4 mM in 5% FFA-free BSA in PBS and incubated at 37°C for 10 min. The fatty acid-BSA solution was then mixed 1:9 with serum-free DMEM with 1% penicillin-streptomycin to obtain a 400 μM fatty acid, 0.5% BSA solution in DMEM (41, 42).

For albumin measurements, media was collected at the end of 48 h of lipid loading and albumin concentration was measured with a commercially available human albumin ELISA kit (Invitrogen, Waltham, MA).

After 48 h lipid loading, media was changed to DMEM with 1% penicillin-streptomycin with or without the addition of cytoskeletal drugs: either 5 μM blebbistatin (MilliporeSigma), 5 μM latrunculin (MilliporeSigma), or 10 μM nocodazole (Cayman Chemicals, Ann Arbor, MI). Cells were treated for 4 h before fixation.

### Traction Force Microscopy

To measure cell traction forces, PHH were seeded on stiff PAA gels embedded with 1 μl/mL red fluorescent beads (Fluorospheres, Invitrogen). Cells were cultured and treated with lipid as described. Brightfield images of cells on the gels and fluorescent images of the beads were taken and the locations marked on a Zeiss Widefield microscope. Cells were dissociated with the addition of 1% Triton-X 100 for 10 min. Fluorescent images were taken again to determine the bead location without the cells attached. Bead images from before and after cell detachment were aligned using the Linear Stack Alignment with SIFT option of the Registration plugin in FIJI. Displacement fields of the beads were generated using the PIV plugin and then further analyzed with the FFTC plugin to calculate forces. Cell outlines were traced from the brightfield images and used to define larger regions of interest (ROIs). Automatic thresholding was used to isolate the areas where traction forces were being generated in proximity to the cell boundary. These thresholded ROIs were used to measure mean and maximum traction forces.

### Immunofluorescence Staining and Microscopy

Following culture, cells were fixed with 4% paraformaldehyde (Thermo Fisher Scientific) for 15 min and permeabilized with 0.1% Triton X-100 in PBS for 15 min. For lamin A/C and HNF4α staining, the permeabilization step was increased to 0.5% Triton X-100 in PBS. Samples were blocked with 5% normal goat serum (Jackson ImmunoResearch Laboratories, West Grove, PA) in PBS for 1 h. Primary antibody for α-tubulin (1:1000, T6199 MilliporeSigma), lamin A/C (1:200, sc-7292, Santa Cruz), or γH2AX (1:100, 05-636-25UG, MilliporeSigma) was applied for 2 h at RT followed by secondary antibody (fluorophore-conjugated IgG, 1:500; Jackson ImmunoResearch Laboratories) for 1 h at RT. Pre-conjugated Alexafluor-647 HNF4α (1:100, ab217073, abcam) was applied for 2 h at RT.

To stain neutral lipids, cells were incubated with BODIPY (1:1,000; Thermo Fisher Scientific) for 1 h at 37°C. Actin was stained with phalloidin (1:50; Invitrogen) for 30 min at RT and nuclei with DAPI (1:5,000; Thermo Fisher Scientific) for 15 min at RT. Samples were washed with PBS between steps. Samples were then mounted with aqueous mounting medium (KPL, Gaithersburg, MD), and 40x images were taken using a Leica TCS SP8 laser scanning confocal microscope. An additional 10x digital magnification was used to zoom in on individual nuclei for measurements of irregularity, γH2AX foci, and lamin A/C and HNF4α intensity.

### Image Analysis

Fluorescence intensity of various proteins was analyzed with FIJI. Fluorescence intensity and integrated density were quantified within individual cells manually segmented by tracing cell outlines. Nuclei were segmented by smoothing and thresholding the DAPI-stained regions and then filling the holes of the binary image. Segmented nuclei were then postprocessed in MATLAB to quantify the nuclear irregularity or used as ROIs within FIJI to measure nuclear area and HNF4α intensity. γH2AX foci within the nucleus were counted manually in FIJI.

Cell volumes were calculated using the 3D Imaging Toolbox for FIJI on confocal Z-stacks of phalloidin staining. Reconstruction of confocal z-stacks is a common method for estimating cell volume (43, 44). The cell boundary was segmented by applying smoothing and by thresholding the phalloidin stain in each slice. The binary images were processed to fill holes to generate an ROI inclusive of the entire cytoplasmic volume. Individual cells were then segmented using the Simple Segmentation Tool applied to the thresholded stack and cell volumes were calculated using the 3D ROI Manager. Nuclear volumes were calculated similarly with the DAPI channel used to segment the cell nuclei. Specifically for the cell vs nuclear volume and cytoplasmic vs nuclear volume plots, volumes were analyzed using a MATLAB program that would calculate the cell, nuclear, and lipid volume in each cell. This was done first by smoothing and thresholding the phalloidin channel of 3D confocal image stacks, which was used to generate a label matrix to identify individual cells. 3D image properties (regionprops3) were used to calculate cell volume. The cell label matrix was then used to calculate the nuclear and lipid volume (in the DAPI and BODIPY channels respectively) within each cell ROI, applying a mask of each cell and then calculating volume with regionprops3. Cytoplasmic volume was estimated by subtracting nuclear and lipid volume from the cell volume.

Nuclear irregularity was quantified with a custom MATLAB program. Binary images of the cell nuclei were generated in FIJI as described and read into MATLAB. The MATLAB Imaging toolbox was used to identify the nuclear boundary and the center of each nucleus. The program linearized the membrane boundary by measuring the distance between the center point and each point along the boundary and plotted this against the angle (see Figure 2b for schematic). To account for the difference in cell area, the linearized membrane boundary was normalized to the mean radius of the nucleus. The program then calculated the nuclear irregularity: the area between the membrane boundary and a perfect circle with the same mean radius. This parameter is sensitive to both large- and small-scale nuclear deformations as both will sum together in generating this irregularity parameter. The linearized membrane boundary can also be further analyzed to find the local curvature. The membrane boundary is smoothed with a loess filter and points of inflection are estimated using two finite differences (analogous to taking the second derivative of a continuous function). Segments of the membrane boundary between two inflection points were then fit with a circle using the Pratt method (45). The radii of curvature along the membrane was collected for each cell and pooled within a group and the probability distribution was estimated with a histogram. Nuclear YZ Aspect Ratio was calculated by re-slicing confocal z-stacks, taking the maximum projection and thresholding to segment each nucleus.

Chromatin condensation analysis was performed according to a previously published method (46). In brief, the raw images were down-sampled and the intensity redistributed before a Sobel edge detection filter was applied. This was followed by automatic thresholding and morphological thinning as well as removal of the nuclear outline. Finally, the chromatin condensation parameter is the number of remaining edge pixels divided by the nuclear area.

Actin and microtubule fiber length and branching density were determined using the RidgeDetection plugin for FIJI. The maximum z-projection of confocal z-stacks was taken and individual cells segmented. The individual cells were then analyzed for mean fiber length, total fiber length, and total number of junctions. The junction density was calculated by dividing the total number of junctions in the cell by the total fiber length. Actin fiber alignment was further analyzed in MATLAB based on the FINE alignment analysis (47). The orientation distribution for each cell was determined with Fourier-based image analysis and then the cumulative orientation distribution was fit with a sum of sigmoid functions. Each sigmoid function represents a fiber family aligned in a specific direction. Cells were then classified based on the number of fiber families detected. Distribution of actin and microtubule density as a function of height was determined by measuring the integrated density of staining in each slice of the stack. The integrated density in each slice was then divided by the total integrated density to determine the normalized distribution from bottom to top of the cell. The normalized distribution was resampled in MATLAB to normalize to the cell height and averaged across all cells, then fit with a smoothing spline to visualize distribution.

### Theoretical Modeling of Chromatin Condensation

To investigate the effect of epigenetic regulation and mechano-osmotic loading on the nucleus, we developed a mathematical model for chromatin phase-separation in the nucleus. The composition of the nucleus at any point and time was defined in terms of volume fractions heterochromatin *ϕ*_*h*_(*x, t*), euchromatin *ϕ*_*e*_(*x, t*) and nucleoplasm *ϕ*_*n*_(*x, t*) such that *ϕ*_*e*_ + *ϕ*_*h*_ + *ϕ*_*n*_ = 1. Equivalently, the physical state of the nucleus at any point can be completely determined via two independent variables – *ϕ*_*n*_, the volume fraction of nucleoplasm, and *ϕ*_*d*_ = *ϕ*_*h*_ − *ϕ*_*e*_, the difference between the volume fractions of heterochromatin and euchromatin. We constructed a free energy density function written as,

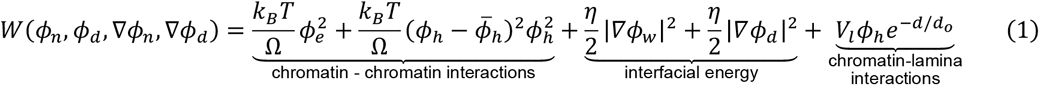

The first two terms in Eq. (1) denote the energetic contributions arising from the competition between entropy and enthalpy of mixing of the two distinct phases – euchromatin phase with *ϕ*_*h*_ = 0 and heterochromatin phase with 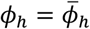. The second term denotes the interfacial energies penalizing the formation of interfaces between the two phases, while the last term for *V*_*l*_ <0 captures the effect of proteins such as LAP2*β*, and LBR which mediate the interactions between chromatin and the nuclear lamina. Note that the chromatin-lamina interactions decrease with distance *d* from the lamina, over a length scale *d*_0_.

The steady state chromatin organization in the nucleus was obtained as local minima of the total free energy defined using Eq. (1), giving rise to the governing equations of the steady state using variational principles as,

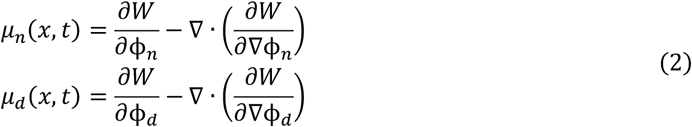

where *μ*_*n*_ and *μ*_*d*_ are the chemical potentials of nucleoplasm and chromatin, respectively. Spatial gradients of the chemical potential of reactively inert nucleoplasm drive its spatio-temporal evolution via the diffusion kinetics as,

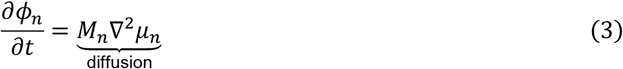

where *M*_*n*_ is the mobility of nucleoplasm in the nucleus. The kinetics of chromatin evolution is driven by both the diffusion as well as epigenetic regulated reaction kinetics of acetylation and methylation as,

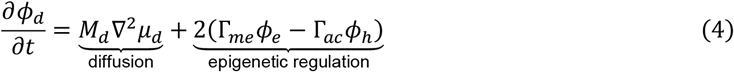

where *M*_*d*_ is the mobility of chromatin in the nucleus and Γ_*me*_ and Γ_*ac*_ are the effective rates of methylation and acetylation of chromatin, respectively. Eq. (2), (3), and (4) together govern the organization of chromatin in the nucleus.

In addition, the nuclear envelope may enforce an interchange of water with the cytoplasm via water exchanging channels enforcing a chemical potential 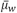 at the nuclear periphery acting as the first boundary condition. Lastly, we assumed a no flux boundary condition along the nuclear periphery for chromatin kinetics. We introduced small perturbations to *ϕ*_*d*_ and *ϕ*_*n*_ around their initial values to initiate the separation of chromatin into the two phases, and let the simulation proceed until a steady state was reached.

### Theoretical Modelling of Cytoskeletal/Lipid Droplet Interactions

To study the effect of lipid droplets on the chemo-mechanical behavior of cells, we used our theoretical cell model previously developed in reference (48). The three-dimensional cell model includes the following components: the cytoskeleton, the focal adhesions, and the nucleus (see reference (44) for details).

The cell cytoskeleton was treated as a continuum of representative volume elements (RVEs), each of which was comprised of (i) the myosin motors, (ii) the microtubules, and (iii) the actin filaments. Myosin molecular motors are the first element of the cytoskeleton in our model and generate internal contractility of the cell as experimentally reported(49). We treated the average density of phosphorylated myosin motors as a symmetric tensor *ρ*_*ij*_, whose components represent cell contractility in different directions(50). The cell contractility *ρ*_*ij*_ generates compressive stress 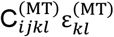 and tensile stress σ_*ij*_ in the cytoskeletal components that are in compression (e.g., microtubules) and tension (e.g., actin elements), respectively,

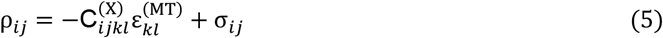

where 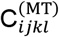 and 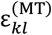 are the stiffness and strain tensors of the cytoskeletal components that are in compression. Actin filaments are the second component of the cytoskeleton which are connected to the myosin element in series and subsequently experience tension and transmit myosin-generated tensile forces to the extracellular matrix through focal adhesions as experimentally observed(51-53). Microtubules are the third component of the cytoskeleton which are connected to the contractile myosin element in parallel and therefore experience compression consistent with experimental observations.

Focal adhesions in our coarse-grained model were modeled as a set of initially soft nonlinear mechanical elements that stiffen with tension to capture the tension-dependent formation of the focal adhesions. When the tensile stress exerted by the contractile cell to the adhesion layer exceeds a certain threshold, mature focal adhesions are formed, and the cell is connected to the substrate, while below this threshold, the stiffness of the adhesion layer remains low and the substrate experiences negligible forces. The nucleus was treated as an elastic thin layer (representing the nuclear envelope) filled with a solid elastic material representing chromatin and other subnuclear components. To perform traction force microscopy simulations, the cell model was coupled to the matrix model which treats the matrix substrate as a thick linear elastic material with an elastic modulus of 10 kPa as used in our traction force microscopy experiments. We treated the lipid droplets as growing mechanical inclusions within the cytoplasm. To this end, we modeled each droplet as a thin spherical membrane with internal pressure representing the enclosed fluid. We simulated the internal pressure by applying uniform and outward force spatially perpendicular to the internal surface of the membrane. As a result of the internal pressure, the memebrane undergoes tensile stresses tangential to the membrane surface representing the surface tension in lipid droplets. In our model, lipid droplets can only have mechanical interactions with other cellular components and the model does not include any chemical effects of lipid droplets on cell behavior. We used a frictionless contact mode in our model for the contact between droplets and the nucleus as well as between droplets and the matrix substrate.

### Statistics and Data Analysis

All data represent experiments using hepatocytes from at least three different biological donors. In each plot, measurements from individual cells are shown in grey to give a sense of the total variation while the average measurements from the biological replicates are shown with the colored dots. To avoid over confidence, statistics were performed on the biological replicates using a two-way analysis of variance (ANOVA) that fit a full-effect model followed by multiple comparisons with Tukey correction. In the case of the drug-treated cells in Figure 6 (where the n number for the drug-treated cells and controls were different), a two-way ANOVA that fit only a main-effects model was used, followed by a Fisher’s test for multiple comparisons. Correlations were analyzed with linear regression followed by an F-test to determine whether the slope was significantly different from zero. Analysis of covariance was then also run to compare the slope and intercept of the fit between two groups. When slopes and intercepts were not significantly different, a pooled slope and intercept could be generated and fit to the pooled data. The distribution of indent radii was compared between control and oleate treated cells using multiple Kolmogorov–Smirnov (K-S) tests, one for each stiffness. The symmetry of indent distributions was determined by comparing the positive dent distribution with the absolute value of the negative dent distribution with a K-S test for each group. The frequency of γH2AX-positive nuclei was compared using multiple Chi-Squared tests comparing control to oleate-treated cells within a stiffness. Comparison of nuclear area and irregularity of fully γH2AX-positive nuclei was compared with an unpaired t-test. The prevalence of different numbers of actin fiber families was compared using multiple Chi-Squared tests comparing control to oleate-treated cells. Traction force measurements were compared using a two-way ANOVA with a full effect model on the biological replicates and the shown p-value is the column factor. For all graphs, significance values less than p = 0.05 are labeled. Color-coded significance bars indicate stiffness-dependent differences within a treatment group, either blue for BSA or green for oleate.

## Acknowledgments

We are grateful to the UPenn Cell and Developmental Biology Microscopy Core and the UPenn NIDDK Center for Molecular Studies in Digestive and Liver Disease (NIH-P30-DK050306) for their assistance in providing training and access to the microscopes used in this study. This work was supported by the following funding sources: National Institute of Health grant U54 CA193417 (AEL, RGW); Center for Engineering MechanoBiology, National Science Foundation Science and Technology Center CMMI: 15-48571 (AEL, FA, AK, PAJ, VBS, RGW); National Science Foundation Graduate Research Fellowship Program grant DGE-1845298 (AEL)

## SUPPLEMENTAL MATERIAL

**Fig. S1.**
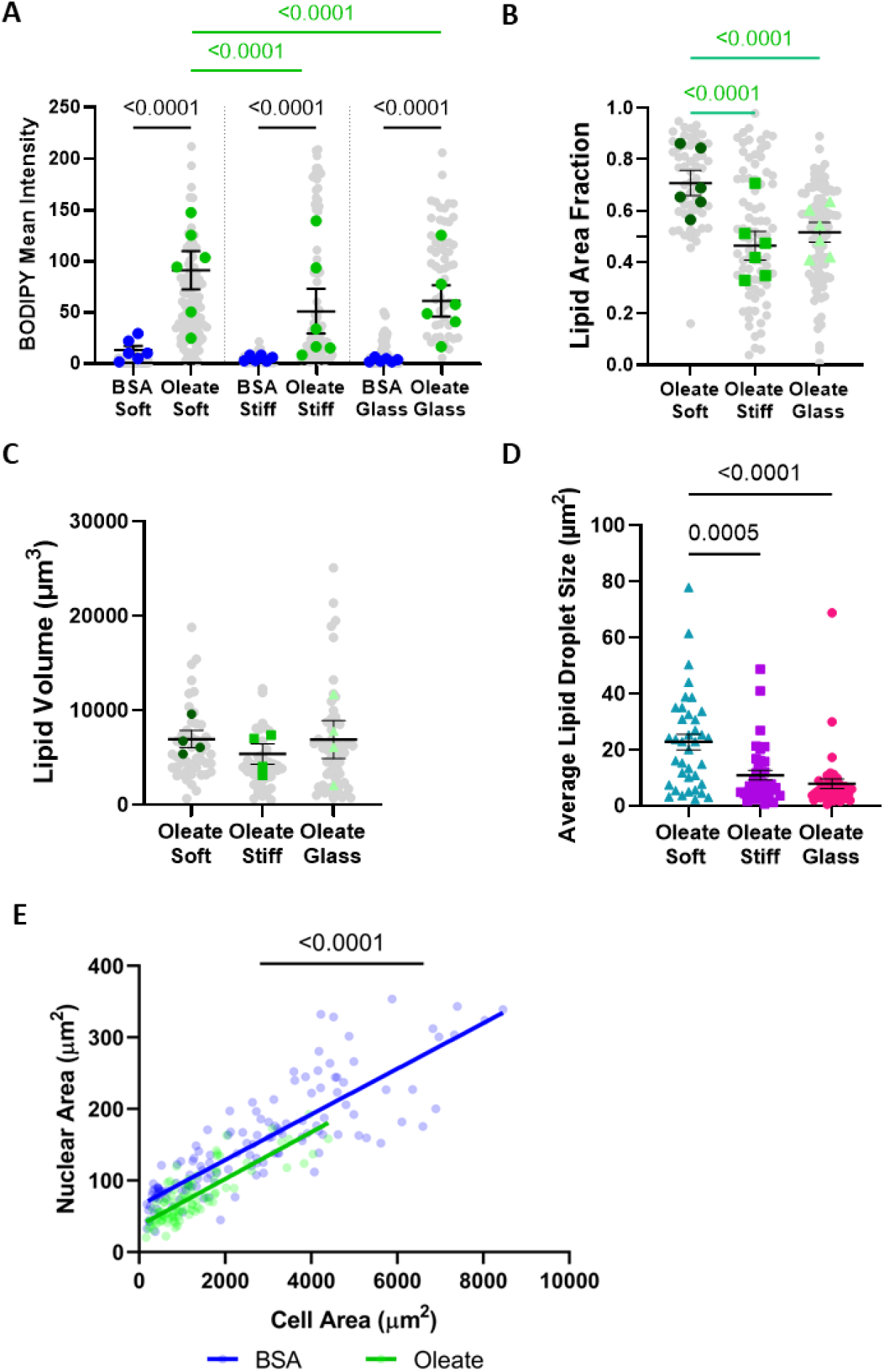
Oleate-treated cells accumulate significant amounts of lipid on all stiffness substrates. (**A**) BODIPY mean intensity for control and oleate-treated cells on soft PAA, stiff PAA, and glass as measured in maximum z-projections. (**B**) Lipid area fraction (lipid positive area/cell area) in oleate-treated cells on different stiffnesses as measured in maximum z-projections. (**C**) Lipid volume in oleate-treated cells on different stiffnesses as measured with 3D segmentation of z-stacks. (**D**) Average lipid droplet cross sectional area per cell of oleate-treated cells from n = 3 independent experiments. (**E**) Scatter plot comparing cell area and nuclear area in individual cells, pooled across stiffnesses, fit with linear regression (n = 3). Statistics: (A, B) Cells are from n = 6 independent experiments. (C, D) Cells are from n = 4 independent experiments. P-values calculated with two-way ANOVA with multiple comparisons. (A-C) Colored dots are the means per experiment, while the grey dots are the individual cell values. (E) Data are the values for individual cells from *n = 3* independent experiments. Blue indicates control cells, while green indicates oleate-treated cells. Slopes of both lines are significantly non-zero (P=<0.0001) but are not significantly different from one another. Y-intercepts are significantly different (P=<0.0001).

**Fig. S2.**
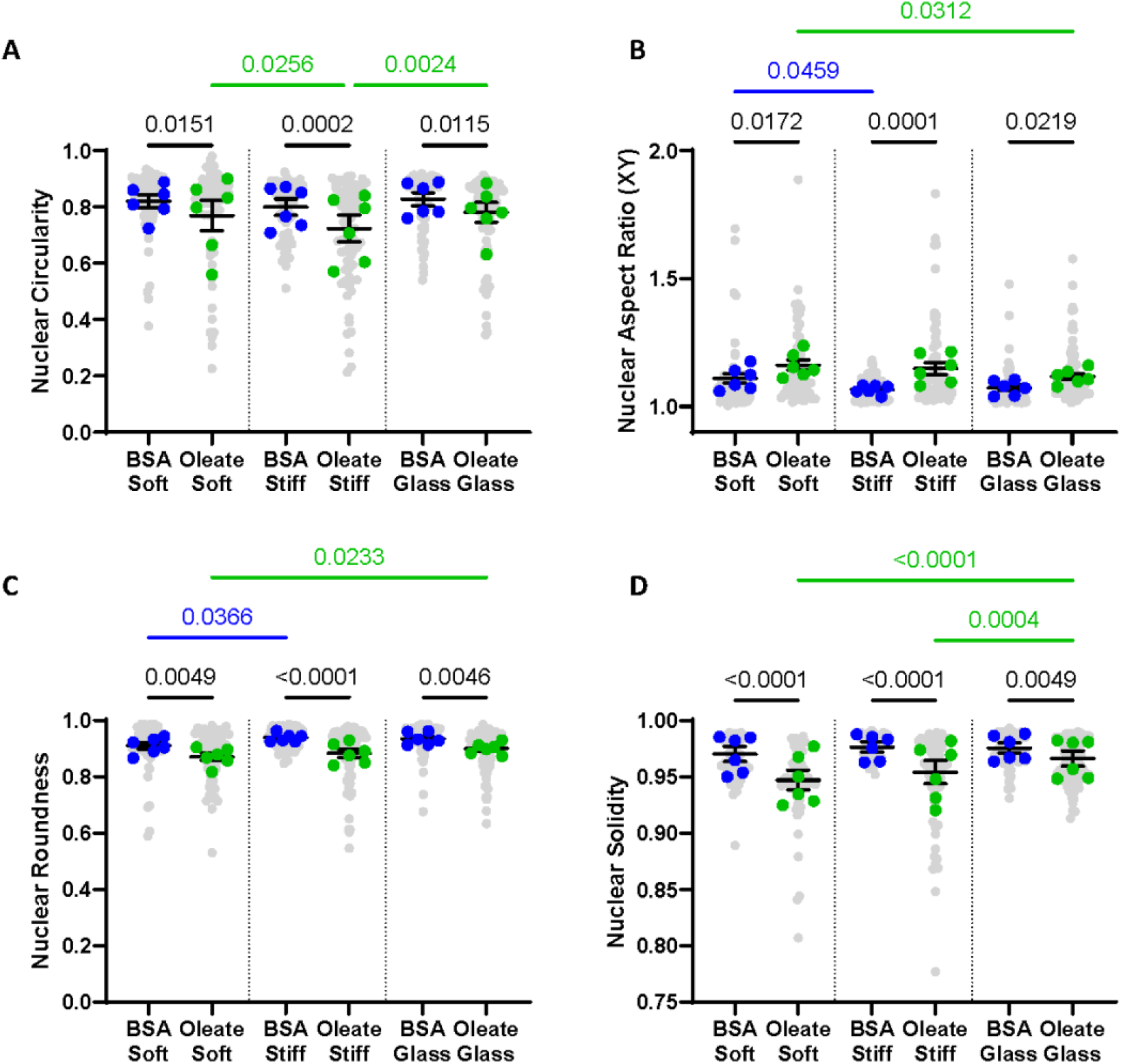
Nuclei of oleate-treated cells are highly deformed according to multiple shape parameters. (**A**) Nuclear circularity, (**B**) aspect ratio (in the XY plane), (**C**) roundness and (**D**) solidity in control and oleate-treated cells on different stiffness substrates. Cells from *n = 6* independent experiments, p-values calculated by two-way ANOVA with multiple comparisons.

**Fig. S3.**
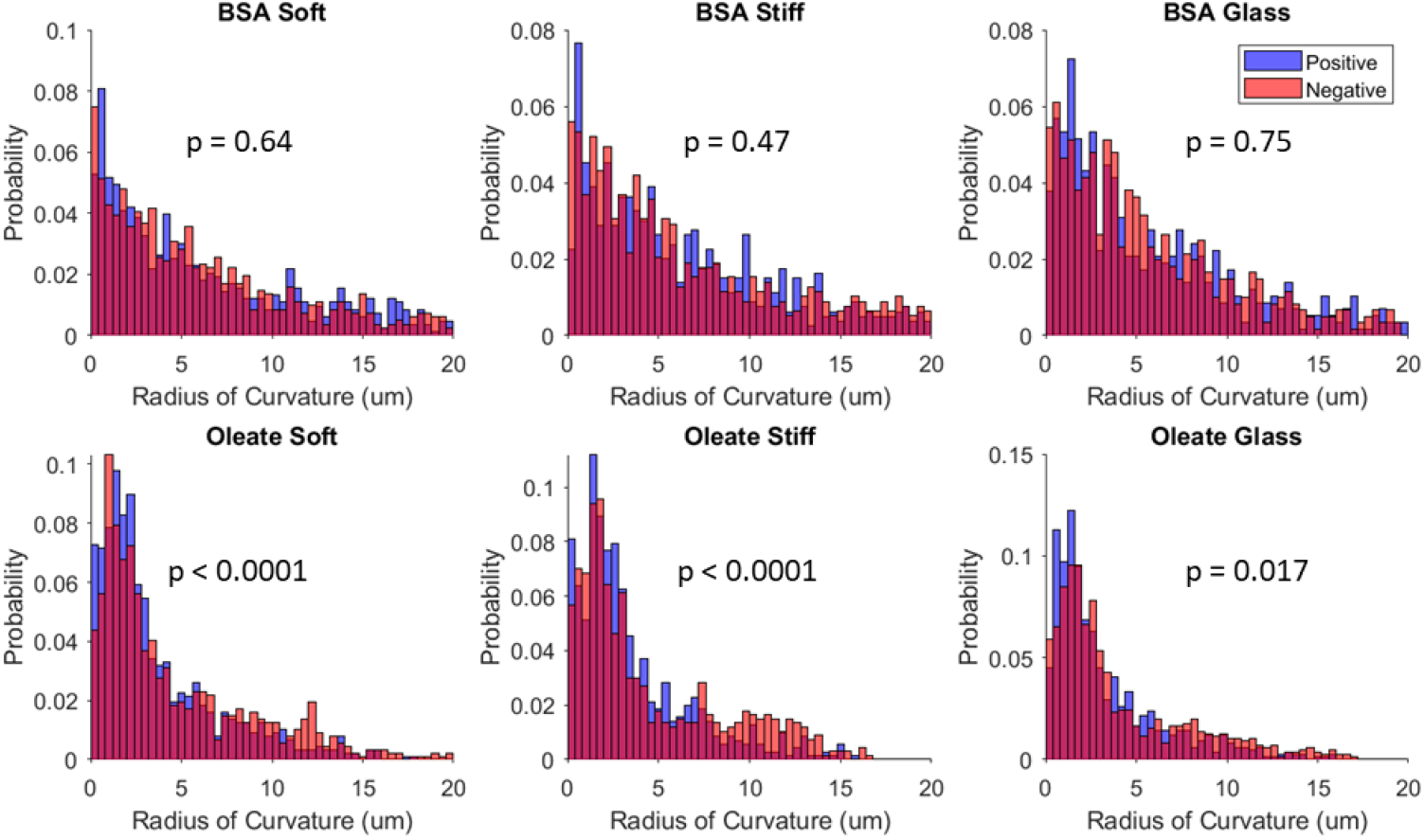
Distribution of radii of curvature for nuclear membrane indentations is symmetric for control but not oleate-treated cells. Overlaid histograms of the magnitude of positive (blue) and negative (orange) radii of curvature for control and oleate-treated cells on soft PAA, stiff PAA, and glass. Radii are all the indents from individual cells in *n = 3* independent experiments. P-values calculated with K-S tests showing oleate-treated cells do not have symmetric distribution (distribution of positive and negative radii is not the same).

**Fig. S4.**
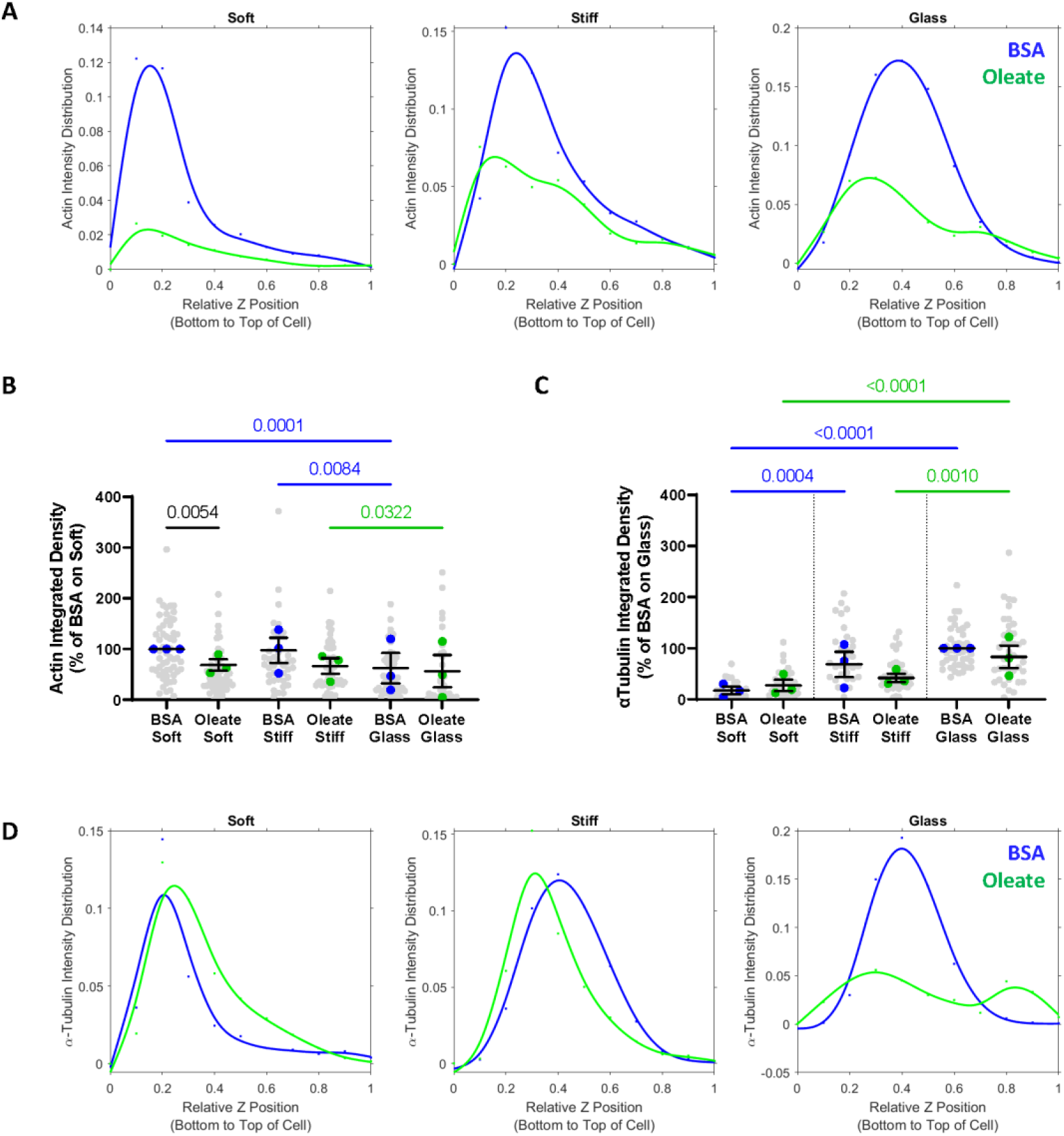
Lipid droplets alter the apical-basal distribution of cytoskeletal fibers. (**A**) Distribution of phalloidin integrated density from the basal to apical membrane of control and oleate-treated cells on soft PAA, stiff PAA, and glass, fit with a smoothing-spline. (**B**) Phalloidin integrated density, indicating total actin content, in control and oleate-treated cells. (**C**) α-tubulin integrated density, indicating total microtubule content, in control and oleate-treated cells. (**D**) Distribution of alpha-tubulin integrated density from the basal to apical membrane of control and oleate-treated cells on soft PAA, stiff PAA, and glass, fit with a smoothing-spline. Cells for all panels from *n = 3* independent experiments, p-values calculated by two-way ANOVA with multiple comparisons.

**Fig. S5.**
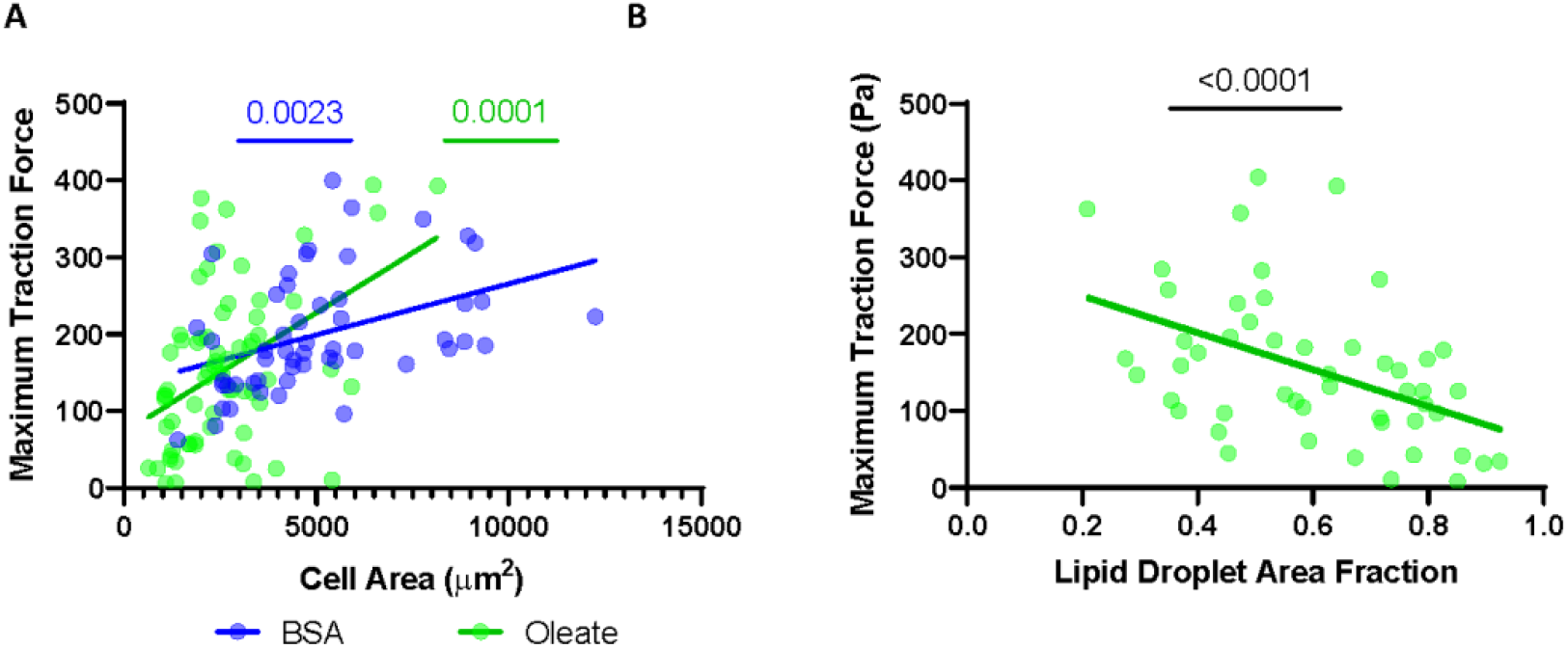
Maximum traction forces decrease proportional to lipid-loading. (**A**) Scatter plot with linear regression of cell area versus maximum traction force in individual cells for control and oleate-treated cells. (B) Scatter plot and linear regression of lipid droplet density and maximum traction force in oleate-treated cells. Data are individual cells from *n = 3* independent experiments. P-values calculated with F-test.

**Fig. S6.**
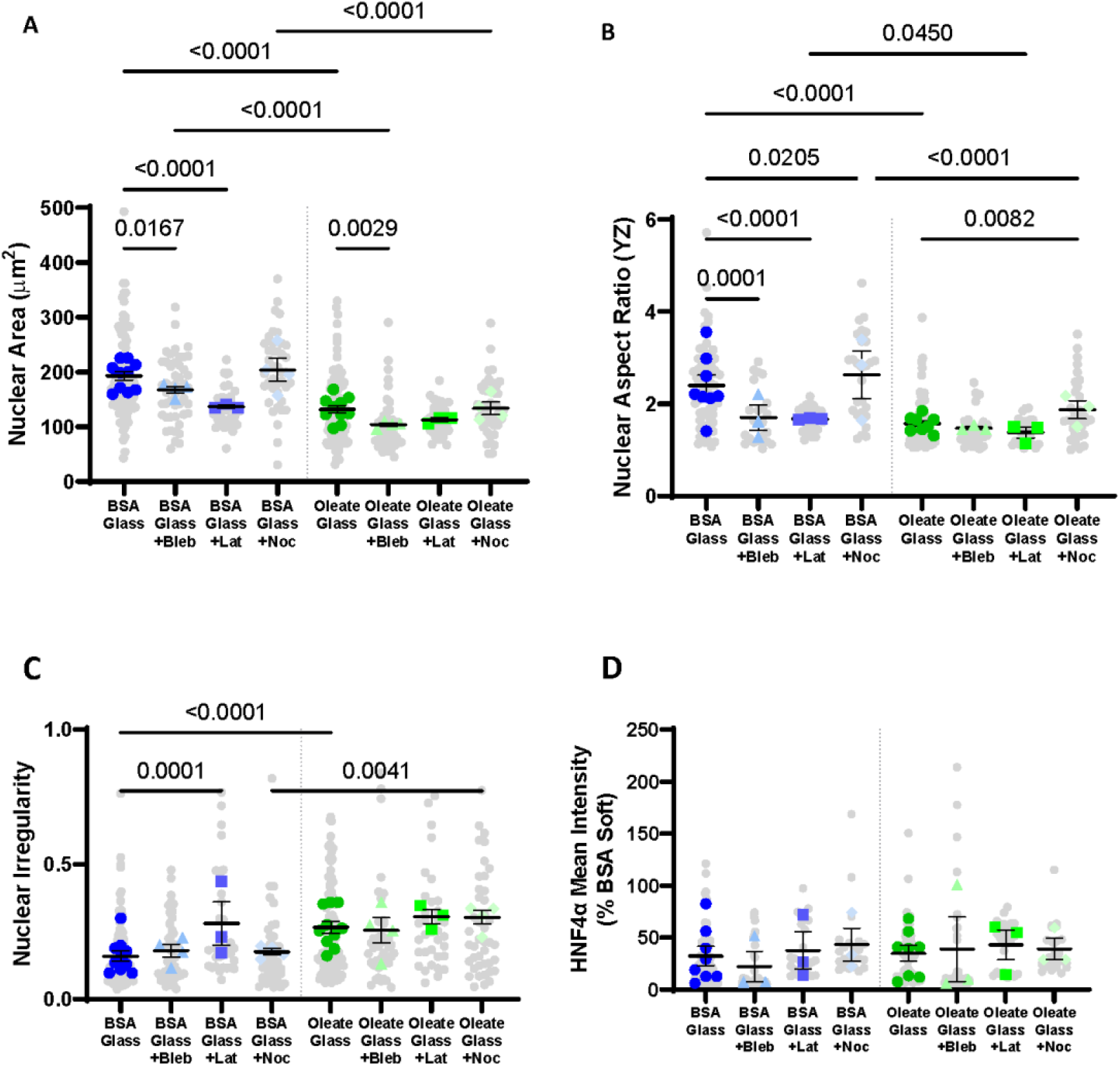
Control cells on glass respond as expected to cytoskeletal drugs. (**A**) Nuclear area, (**B**) cross-sectional nuclear aspect ratio, (**C**) nuclear irregularity, and (**D**) mean HNF4α intensity (normalized to control on soft) in control and oleate-treated cells with or without the addition of cytoskeletal drugs on glass. Blebbistatin (Bleb), Latrunculin A (Lat), Nocodazole (Noc). Data are the mean +-s.e. of *n = 3* independent experiments for each drug treatment and *n = 8* for the non-treated. P-values were calculated using two-way ANOVA with a main effects model with multiple comparisons.

